# Probing tissue-scale deformation by *in vivo* force application reveals a fast tissue softening during early embryogenesis

**DOI:** 10.1101/167155

**Authors:** Arturo D’Angelo, Kai Dierkes, Carlo Carolis, Guillaume Salbreux, Jérôme Solon

## Abstract

During development, cell-generated forces induce tissue-scale deformations to shape the organism. Here, we present a method that allows to quantitatively relate such tissue-scale deformations to spatially localized forces and measure mechanical properties of epithelia *in vivo*. Our approach is based on the application of controlled forces on microparticles embedded in individual cells of an embryo. Combining measurements of the bead displacement with the analysis of induced deformation fields in a continuum mechanics framework, we can quantify tissue material properties and follow their change over time. In particular, we uncover a rapid change in tissue response occurring during *Drosophila* cellularization, resulting from a softening of the blastoderm and an increase of external friction. Pharmacological treatments reveal that in addition to actomyosin, the microtubule cytoskeleton is a major contributor to epithelial mechanics at that stage. Overall, our method allows for measuring essential mechanical parameters governing tissue-scale deformations and flows occurring during morphogenesis.

## Introduction

During animal development, cell-generated forces, which are modulated in space and time, are translated into tissue-scale deformations that shape the living organism (Collinet et al., 2015; Lye et al., 2015; Petridou et al., 2017; Rauzi et al., 2015). The extent and pattern of these deformations depends not solely on the temporal and spatial profile of the generated force fields but also on the mechanical properties of tissues that the force acts on. For instance, when a thin sheet of elastic material moves with friction relative to an external substrate, the range of deformation triggered by an external force is larger in a stiffer elastic material. It is thus conceivable that, similar to cell-generated forces, the mechanical properties of tissues are modulated during development in order to drive morphogenesis towards specific developmental endpoints. However, the direct assessment of the material properties governing tissue-scale deformation in living organism is challenging, as it requires the application of controlled ectopic forces and the concurrent measurement of epithelial deformation within developing embryos. Recent approaches have started to develop methodologies to apply controlled forces employing tissue-embedded (ferromagnetic) fluid droplets combined with force application by means of a magnetic field or by trapping cell contacts with optical tweezers (Bambardekar et al., 2015; Campas et al., 2014; Serwane et al., 2016). Due to the applied force amplitudes, however, droplet movements as well as tissue deformations reached with these experimental techniques remain highly localized, thus hindering the accessibility of mechanical properties emerging on the tissue scale. Here, we present a method overcoming this limitation. In the spirit of previous *in vitro* methods, we make use of a micron-sized functionalized magnetic particle that is pulled on by means of a custom made electromagnet (Bausch et al., 1998; Kollmannsberger and Fabry, 2007). Due to the high force amplitudes available within our assay, large bead displacements and, in particular, large-scale tissue deformations can be induced. We show that quantifying the tissue response to force application on the bead allows to characterize key tissue mechanical properties. Importantly, we do not only monitor bead displacement, but also quantify tissue deformation based on image-segmentation data. By analyzing the deformation field within a continuum mechanics framework, quantitative insights into global mechanical tissue properties can be obtained. In particular, we show that a rapid change in tissue mechanics occurs during cellularization on time-scales of as little as several minutes. Using drug treatments, we find that in addition to the actomyosin cytoskeleton, microtubules also contribute to the material properties of the syncytial blastoderm. Finally, we detect changes in the friction coefficient that correlate with changes in the distance between the blastoderm and the vitellin envelope, indicating that friction is modulated on short time scales during development.

## Results

### Magnetic particle injection and positioning

We have developed a protocol for injecting an individual magnetic microparticle into a living *Drosophila* embryo and for a consecutive embedding of the particle into a tissue of interest: (i) a magnetic bead with a diameter of 4.5 *μm* is injected into the embryo at preblastoderm stage, (ii) the bead is then directed with a permanent magnet towards the surface of the embryo (Figure 1A). The bead is afterwards encapsulated within an individual cell while cellularization occurs (Figure 1A and see Material and methods). The cell accommodates to the embedded bead and the organism develops normally until larval stage (Video 1). By choosing the orientation of the embryo during the steering phase, we can ensure the bead to be encapsulated in predefined groups of cells that will develop into specific tissues at a later stage. In particular, we were able to incorporate magnetic beads in the blastoderm at early stages (Figure 1B) and in the epidermis and amnioserosa at later stages (Figure 1-figure supplement 1C). To specifically target intracellular compartments, the microparticle was coated with a GFP nanobody (Kubala et al., 2010), which specifically binds GFP-tagged proteins. In particular, in order to attach the bead to the cell surface, we made use of fly lines expressing GFP-tagged versions of Resille, a membrane protein (Morin et al., 2001).

**Figure 1.**
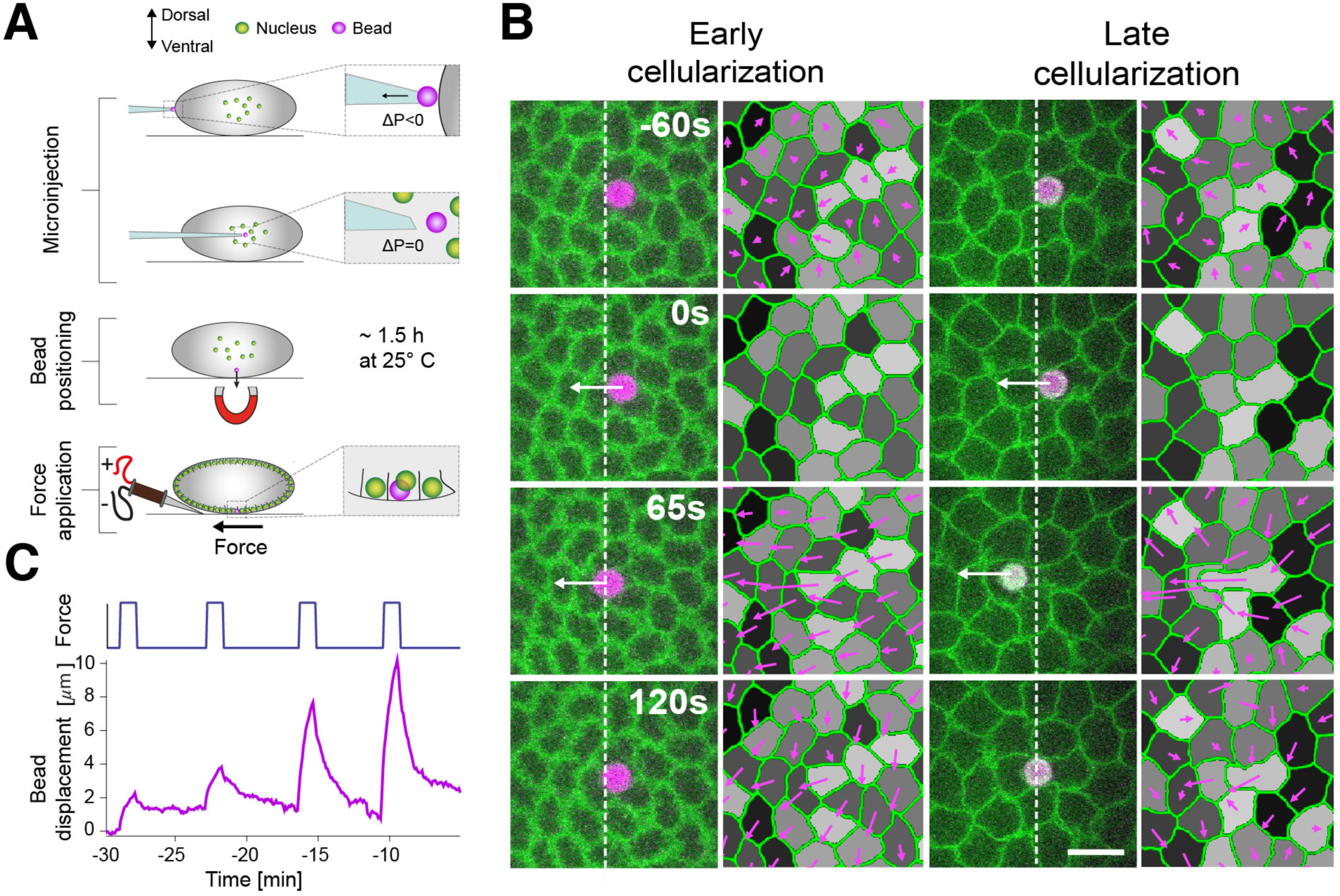
Bead injection and force application. (A) Injection procedure. An individual magnetic bead (purple) of 4.5 *μm* diameter is injected into the yolk of an embryo at developmental stage 2. In order to position the bead apically, the embryo is staged on top of a permanent magnet post injection (~1.5h at 25°C). When cellularization begins, force steps of 65s duration are applied to the bead with an electromagnet. (B) Time lapse images showing bead displacement and tissue deformation (purple arrows) in response to a force step (high force condition, onset at 0s). Here, the bead was embedded into an individual cell of a Resille-GFP embryo. Force applications are shown for early cellularization (left, >20 min before gastrulation) and late cellularization (right, <20min before gastrulation). White arrows indicate force application. White dashed lines mark the left side of the bead at time 0s. Deformations are calculated relative to the onset of force applications (t=0s). Scale bar is 10 μm. (C) Bead displacement for four consecutive force applications at high force condition performed over the time-course of cellularization (bottom) and corresponding forcing curve over time (top). Time is relative to the onset of gastrulation at t=0min.

### Bead displacement and deformation field measurements upon force application

After calibration (see Material and methods), we applied a controlled force step of 65s duration and amplitudes taking values around 115pN to the magnetic bead by means of an electromagnet (Figure 1C, Figure1-figure supplement 1A-B, Suppl. Info. and Material and methods). We obtained two complementary readouts in order to characterize the mechanical response of the tissue: (i) the bead displacement over time and (ii) the deformation field of the apical surface area of the epithelium (Figure 1B-C, see Material and methods).

Applying forces at consecutive times in the blastoderm, spanning a period of 50 min before gastrulation of Resille-GFP expressing embryos, we can observe significant changes in both the displacement of the bead and the induced deformation field (Figure 1B-C, Figure 1-figure supplement 1D, video 2 and video 3). Defining the origin of time as the onset of gastrulation, we find that the amplitude of bead displacement, i.e. the maximal displacement of the bead at the end of the force step relative to its position before the force step, changes abruptly from approximately 2 *μm* to 8 *μm* around t=-20 min (Figure 1C). This change in maximal bead displacement is associated with a concurrent change in the spatial profile and range of the deformation field (Figure 1B and Figure1-figure supplement 1-D). Our data therefore show that the same localized force can lead to a significantly different deformation pattern when applied at developmental time points differing only by a few minutes.

### Spring dashpot analysis shows a fast mechanical switch during cellularization

To obtain a first mechanical description of the response of the tissue upon force indentation, we analyzed bead responses by means of an effective rheological model (Figure 2A). We found that the response to a force step could be well captured by fitting a viscoelastic Maxwell-Kelvin-Voigt model to the individual bead displacement curves. In this mechanical circuit, a viscous element, with viscosity coefficient *μ*_1_, acts in parallel with an elastic element, with stiffness coefficient *K*, both operating in series with a viscous element described by a second viscosity coefficient *μ*_2_ (Figure 2A). In such a rheological scheme, the long time behavior is a fluid relaxation occurring on a timescale dictated by the ratio of the viscosity parameter *μ*_2_ to the stiffness coefficient *K*. Introducing a long-time viscous response was necessary to account for the incomplete relaxation of the bead after the application of the force (Figure 2A), which indicates that the tissue does not behave like a purely elastic material on long time scales.

**Figure 2.**
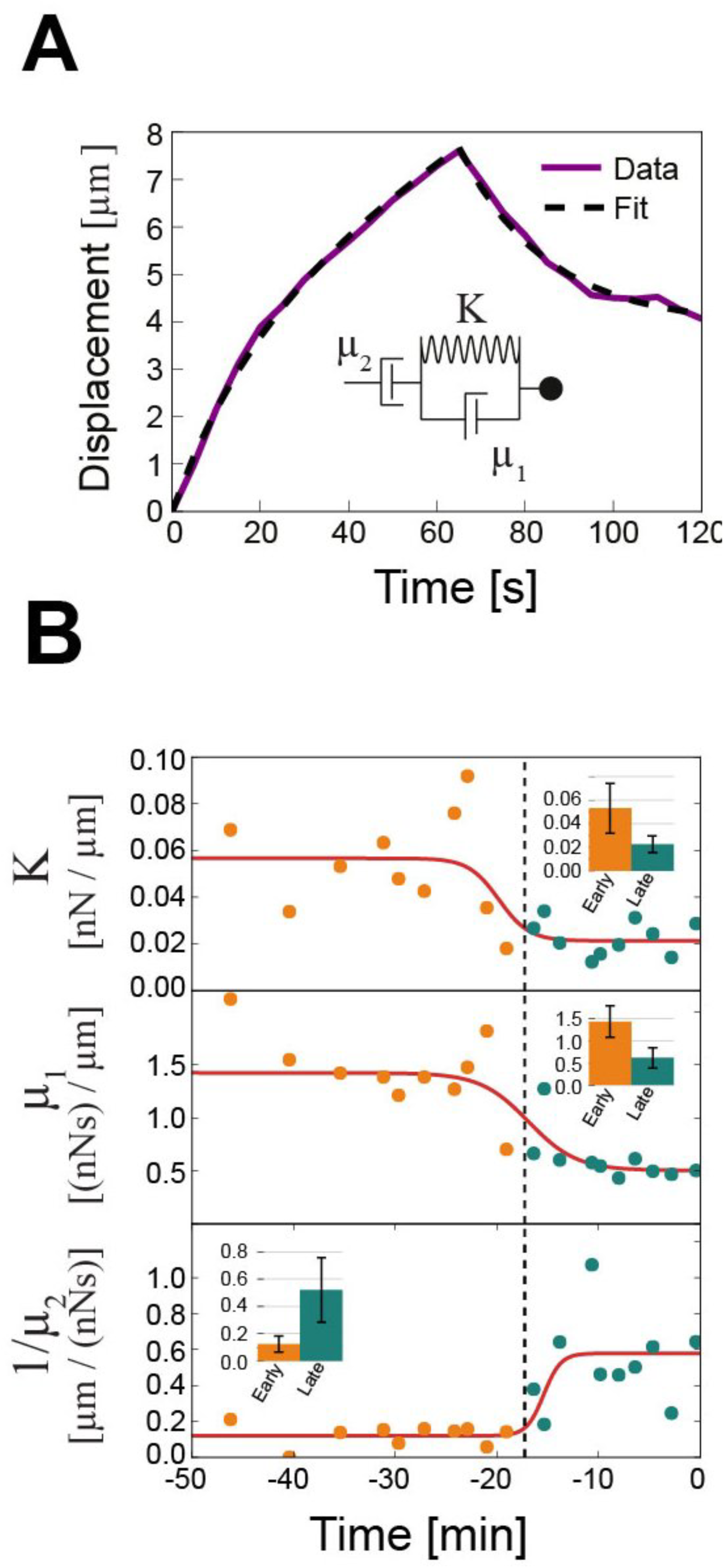
Fitting bead displacement to an effective spring-dashpot model. (A) Example of a bead displacement curve after the application of force (high force condition, purple line) fitted with a Maxwell-Kelvin-Voigt model described by the three effective parameters *μ*_1_(0.61 (nN.s)/μm), *μ*_2_(1.99 (nN.s)/μm) and *k* (0.03 nN/μm) (see inset for a schematic of the corresponding rheological model). (B) Effective parameters as a function of developmental time relative to the onset of gastrulation. Each dot represents a single force application at high force condition. We observe a step-like change of the three parameters at the onset of the fast phase of cellularization, i.e. around -20 min. Red lines are fits of experimental data to a sigmoid function. Black dashed lines represent the mean of the middle of sigmoid curves and is used to separate between force applications in the early (orange) and late (blue) phase of cellularization. Insets are average parameters over the early and late phases, with error bar indicating standard deviations.

We determined the effective parameters introduced above for 20 force application experiments, performed at successive time points during cellularization, the process during which cellular membranes extend basally towards the interior part of the embryo(Figard et al., 2013; Lecuit and Wieschaus, 2000; Royou et al., 2004) (Figure 2B). Plotting them as a function of developmental time reveals a rapid step-like change of all three mechanical parameters, occurring at -17 min before the onset of gastrulation. Indeed, we found that the observed time-courses could be well fitted by sigmoids (Figure 2B). The effective viscoelastic timescales, *μ*_1_/*K* and *μ*_2_/*K*, in contrast, did not change significantly (Figure 2-figure supplement 1C). This observation indicates that both elasticity and viscosities change in the same extent, suggesting that these two parameters are not independent from each other. The observed switch in mechanical parameters coincides with a change in the velocity of progression of cellularization(Figard et al., 2013) (Figure 1-figure supplement 1E-F). We therefore refer to the phases before and after the change in mechanical parameters as early and late cellularization (Figure 1B, Figure 1-figure supplement 1D and Figure 2-figure supplement 1A).

### Identification of changes in intrinsic tissue mechanical parameters and external friction with a continuous 2D viscoelastic description

The analysis of observed bead responses with an effective rheological scheme allows for the detection of rapid changes in the mechanical environment the bead is embedded in. However, effective response parameters are not direct read-Bouts of actual material properties of the tissue, and cannot relate the spatial deformation of the tissue to the applied force. To overcome this limitation, we analyzed two complementary measurements of the tissue response in pulling experiments (bead displacement and tissue deformation) in the framework of a continuum description of tissue mechanics (Figure 3A). In our description, we represent the tissue as a two-dimensional flat sheet with tension tensor *t_ij_*, moving with friction relative to a substrate and subjected to a 2D effective external force density from the bead *f_i_*, such that force balance on the tissue reads

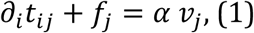

**Figure 3.**
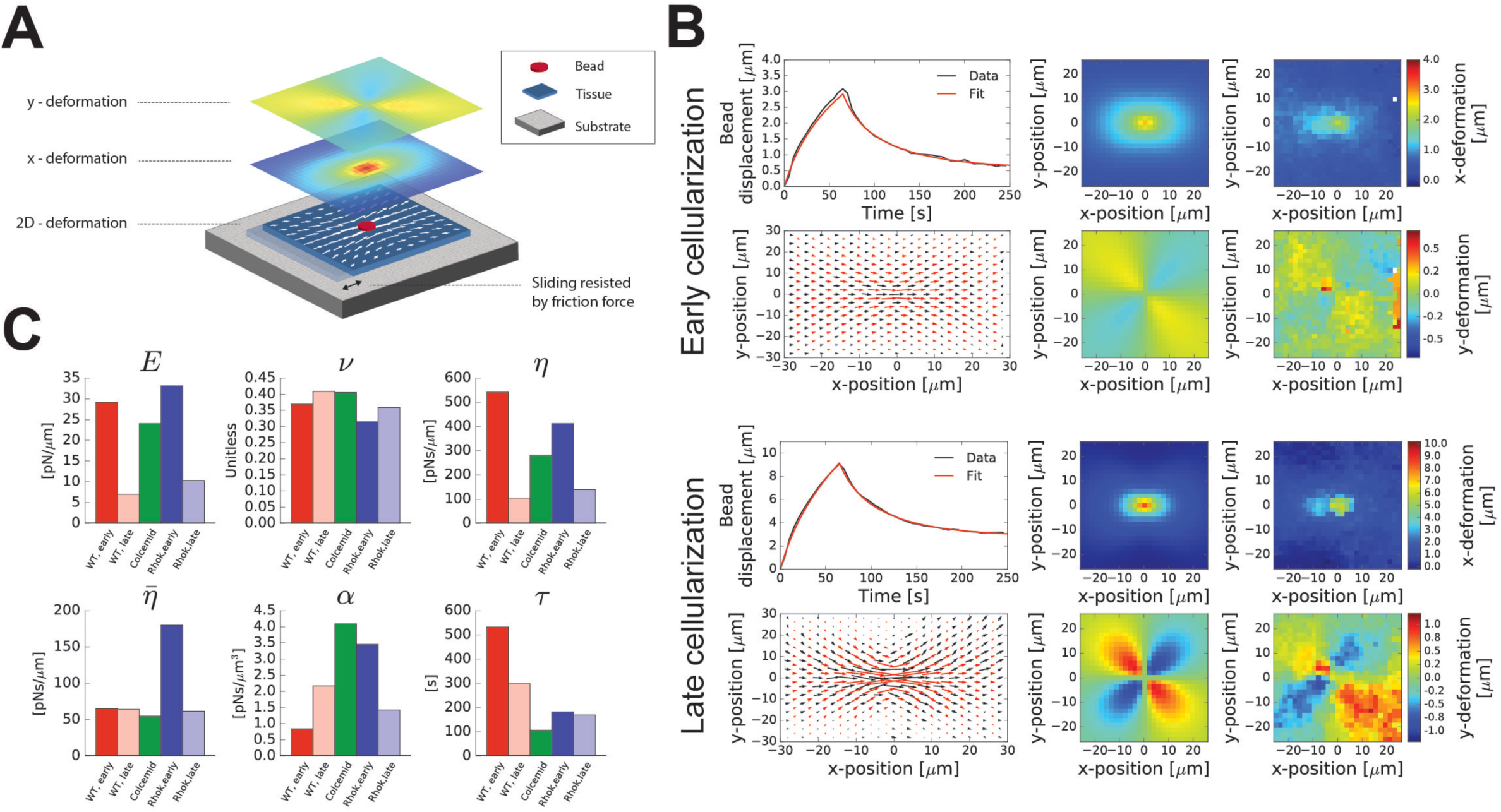
Determining mechanical tissue parameters within a 2D continuum mechanics framework. (A) Schematic of the continuum description used to calculate deformation fields. The epithelial tissue is considered to be a 2D sheet moving relative to a substrate. Sliding of the tissue is resisted by friction. Bead-induced tissue deformation (represented by arrows) is decomposed in x- and y-deformation (shown as heatmaps). Mechanical tissue parameters are extracted from experimental data by fitting the continuum description to both the averaged time-evolution of bead displacement and deformation field at the end of the forcing step. (B) Comparison of experimental and continuum description of bead displacement at early (n=10) and late cellularization (n=10) and deformation field at early (n=10) and late cellularization (n=8). For both conditions, average experimental and fitted bead displacement (top-left panel), fitted (red arrows) and experimental (black arrows) deformation field (bottom-left panel), and fitted and experimental x- and y-deformations (middle and right panels, respectively) are shown. (C) Mechanical parameters as extracted from the fit of experimental data to our continuum description. Values of the elasticity *E*, the poisson ratio *v*, the shear viscosity *η*, the bulk viscosity 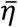, the friction coefficient α, and the Maxwell viscoelastic timescale τ are shown for both WT and embryos injected with Colcemid and Rho-K inhibitor.

where *α* is a friction coefficient and *v* the velocity vector. In addition, we used the following constitutive equation for the tension tensor:

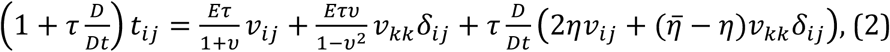

where 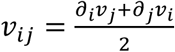 is the velocity gradient tensor, *E* and *v* are a two-dimensional Young’s modulus and Poisson ratio, *η* and 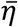 are a shear and bulk viscosity, and τ is a viscoelastic relaxation time. The tissue is described here as a linear Maxwell-Kelvin-Voigt material: the tissue is assumed to have a viscous response characterized by *η* and on short time scales *t* < *η* /*E*, an elastic response on intermediate time scales 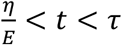, and that elastic stresses are relaxed above the Maxwell viscoelastic time scale *τ*. In line with our choice of a continuous linear description, we note that for forces ranging from 50pN to 115pN, bead displacement and epithelial deformations scale approximately linearly with the force amplitude (Figure 2-figure supplement 1B, Figure 3-figure supplement 1D). Note that Eq. (2) is, however, a description of the tissue rheology which is not equivalent to the effective description of the bead behaviour introduced above, as it distinguishes explicitly between dissipative processes internal to the tissue (characterized by viscosities) and external to the tissue (characterized by a friction coefficient). In addition, Eq. (2) discriminates between isotropic and anisotropic elastic moduli and viscosities.

We then solved Eqs. 1 and 2 and used a fitting procedure to adjust the theoretically predicted bead displacement as a function of time and the displacement field at the end of force application to experimental results (Figure 3B, Suppl. Info. and Material and methods). We found that both the bead displacement and tissue deformation field could be reproduced with the continuum description in the early and late phases of cellularization (Figure 3B, Video 4 and Video 5).

We then compared the values of mechanical parameters extracted at different stages. We found changes in elasticity from *E* ≃ 29 pN/μm to 7 pN/μm, shear viscosity from *η* ≃ 540 pN. s/μm to 105 pN. s/μm, and Maxwell viscoelastic timescale from τ ≃ 530s to 300s (Figure 3C). These changes correspond to an overall softening of the epithelium, while the characteristic timescale of viscous to elastic transition (~10s) and long-time plasticity (~100s) does not vary significantly. Interestingly, these values match relaxation timescales measured during the Zebra fish tail bud (Serwane et al., 2016).

In addition, we found that the friction coefficient was increasing from ~0.8 pN.s/μm^3^ to 2.2 pN.s/μm^3^. We wondered why the friction coefficient was changing over the course of cellularization, given that it reflects resistance to motion between the tissue and surrounding structures. We therefore analyzed the width of the perivitelline space as a function of time, and found that the apical surface of the cells is getting in closer contact with the vitellin envelope at late cellularization (Figure 3-figure supplement 2A-B). This observation suggests that as the embryo approaches the vitellin envelope, increased mechanical interactions lead to an increase in the external friction coefficient.

Finally, we note that the long time-scale hydrodynamic length 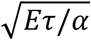 characterizes the spatial range of mechanical interactions in the tissue (Bonnet et al., 2012; Mayer et al., 2010). We find that this length decreases from ~140 μm to ~30 μm over the course of cellularization.

Altogether, our method allowed us to identify a significant tissue softening and an increase in external friction occurring during cellularization. Both are contributing to the strong reduction in the spatial propagation of deformation generated by otherwise similar force fields.

### Actomyosin and microtubule cytoskeletons impact the mechanical properties of the blastoderm

To relate the observed changes in mechanical properties of the epithelium during this softening to intracellular components, we performed our mechanical probing experiments in embryos injected with pharmacological drugs affecting cytoskeletal components. We used a Rho kinase inhibitor (Rho-K), Y27632 (Uehata et al., 1997), and a microtubule depolymerizing drug, Colcemid, to affect actomyosin contractility and the microtubule cytoskeleton, respectively. It has been reported that both components of the cytoskeleton contribute to the developmental remodeling occurring during cellularization (Foe and Alberts, 1983; Royou et al., 2004; Schejter and Wieschaus, 1993; Xue and Sokac, 2016). Pharmacological drugs were injected after magnetic particle injection and before performing pulling experiments. The output of the experiments was then analyzed with both the spring-dashpot model and our 2D continuous viscoelastic model.

Rho-K inhibited embryos did not show major changes in the effective parameters *K, μ*_1_, and *μ*_2_ compared to WT in both the early and late phases of cellularization (Figure 2-figure supplement 1B, 1D, Video 6 and Video 7). A fit to our 2D description indicated that epithelial elasticity is not affected by Rho-K treatment in both phases (Figure 3C and Figure 3-figure supplement 1B-C). However, we observed a major change in bulk viscosity 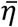 of ~180% at early cellularization, in the friction coefficient *α* compare to WT (a 310% increase at early cellularization and a 34% decrease in late cellularization), and a decrease in the viscoelastic relaxation timescale *τ* (a 66% decrease at early cellularization and a 43% decrease at late cellularization). This is reflected in changes in the spatial propagation of the deformation and of the hydrodynamic length scale by a factor ~3 compared to WT early cellularization (Supp. Info. Table II, Figure 3C, Figure 3-figure supplement 1-B). To identify whether the changes in friction observed in Rho-K injected embryos arise from the interaction with the vitellin envelope, we estimated the width of perivitellin space. We found that the distance between the apical cell surface and the vitelline membrane is smaller in Rho-K treated embryos at early cellularization, i.e. when we measure a higher external friction and, conversely, increased in late cellularization when we measure a decrease in external friction (Figure 3-figure supplement 2A-B).

In the absence of microtubules, apico-basal progression of membranes is impaired and embryos fail to cellularize (Foe and Alberts, 1983; Lecuit and Wieschaus, 2000; Royou et al., 2004). We therefore used Colcemid treatment to measure the effect of microtubule depolymerization on epithelial mechanics during cellularization. We observed that the transition in mechanical parameters observed in WT was absent in Colcemid-treated embryos, indicating that the shift in mechanical parameters is associated with the process of cellularization (Figure 2-figure supplement 1D). Interestingly, Colcemid treatment also affected the response of the epithelium to mechanical forces in the early phases of cellularization. With our spring-dashpot analysis, we identified that the major effect of Colcemid treatment was an increase in the fluidity 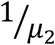 by a factor ~2 compared to WT in the early phase (Figure 2-figure supplement 1B and D). Interestingly, these modulations are also associated with changes in the deformation field. The spatial range of experimentally observed deformation is smaller than in WT at early cellularization (Figure 3B, Figure 3-figure supplement 1B, Video 2 and Video 8). As for WT, we could recapitulate the deformation field and bead displacements measured in Colcemid injected embryos with our 2D theoretical description of the epithelium (Figure 3-figure supplement 1B). Compared to WT in early cellularization, our fitting procedure indicated a 5-fold increase in the friction *α*, a 2-fold decrease of the shear viscosity *η* and a 5-fold decrease in the long-time viscoelastic timescale *τ* (Figure 3C and Figure 3-figure supplement 1B). This decrease in the Maxwell viscoelastic timescale in the continuum model indicates that long time-scale relaxation of elastic forces is considerably accelerated in Colcemid-treated embryos. This observation suggests that relaxation of stresses stored in the microtubule network is responsible for the slow dynamics of tissue deformation in the WT embryo. Using these values, we find that the hydrodynamic length is about 25 μm, i.e. ~5 times shorter than WT in early cellularization, thus accounting for the decrease in the spatial extent of deformation observed in Colcemid-treated embryos. The observed changes in frictional forces are, here as well, consistent with a reduction in the space between the vitellin envelope and the blastoderm cells (Figure 3-figure supplement 2A-B). Altogether, we observe an inverse correlation between friction coefficients and perivitellin space widths (Figure 3-figure supplement 2A-B), suggesting that cytoskeletal components may affect the distance between the tissue and the vitellin membrane, and consequently external friction.

## Discussion

Here, we presented a methodology to apply controlled forces, extract tissue-scale deformations and measure mechanical properties of epithelia within a generic continuum mechanics framework. Our method allows for the *in vivo* measurement of essential physical parameters governing tissue-scale deformations and flows and therefore to relate force patterns to spatial deformation profiles. We can monitor mechanical changes during development, and could uncover a rapid step-like softening of the blastoderm preceding the onset of gastrulation. Our work therefore indicates that, in addition of the modulation of cellular forces, mechanical properties of tissues can change significantly on time-scales of the order of minutes during morphogenesis. Such changes must impact morphogenetic processes, as the pattern and amplitude of deformations in response to the same force field is set by the mechanical properties of the involved tissues. In particular, we can speculate about an essential role for the observed softening during cellularization in the consequent rearrangements associated with gastrulation. Indeed, low tissue stiffness and high external friction reduce the range of propagation of deformations, and therefore allows for local deformation without impacting cells at larger distances. This feature could be important when several morphogenetic rearrangements occur concurrently, such as during gastrulation. In addition, our results obtained in drug-injected embryos identified the microtubule network as one of the main factors in determining tissue mechanics at this stage. Our work therefore suggests that the observed modulation of tissue mechanics is due to the rearrangement of the microtubule network occurring during cellularization (Foe, 1993; Mazumdar and Mazumdar, 2002). Microtubules emanating from the asters associated to different nuclei display transient connections that disappear during cellularization. A highly connected microtubule meshwork before celularization thus rearranges into a network of only weakly interacting microtubules asters. One hypothesis could be that this change in connectivity is at the origin of the observed changes in mechanics.

Finally, our method for mechanical measurements, due to its versatility, low cost and adaptability to different microscopy techniques can easily be employed in other systems. It thus paves the way for further studies mapping out epithelial mechanics. Therefore, it will serve as an important tool for understanding the emergence of mechanical properties at the tissue-scale in developmental contexts and in cases of disease.

## Acknowledgments

We thank Xavier Trepat for discussions and critical reading of the manuscript. We are grateful to the ALMU team for providing help with microscopy. The research leading to these results has received funding from the Spanish Ministry of Economy and Competitiveness, Plan Nacional, BFU2010-16546 and BFU2015-68754 and ‘Centro de Excelencia Severo Ochoa 2013–2017’, SEV-2012-0208. We acknowledge the support of the CERCA Programme/Generalitat de Catalunya. G.S. is supported by the Francis Crick Institute which receives its core funding from Cancer Research UK (FC001317), the UK Medical Research Council (FC001317), and the Wellcome Trust (FC001317).

## Material and methods

### Preparation of *Drosophila* Embryos

To inject the bead at pre-blastoderm stage, the embryos are collected during 15min, dechorionated in 100% bleach and mounted on an heptane/glue coverslip(Fish et al., 2007). The tissue in which the bead will be located has to face the coverslip. Once mounted, the embryos are dehydrated during 10 min at 25°C and then covered with voltalef 10S oil.

### Particle microinjection

MicroBneedles are generated by pulling 1mm glass capillaries (Narishige G1) using a micro-puller (Sutter instruments P30). The needle tip is then opened and beveled in a controlled manner to facilitate the injection using a micro grinder (Narishige EG-44). The internal diameter is set to be slightly smaller than the particle, i.e., 3.5-4 *μm* (Fig 1-A). This allows, by tuning the pressure inside the micro-needle with a microinjector (WPI PV 820), to hold and inject an individual particle within the *Drosophila* embryo. Once inside the embryo, the bead can be oriented on the A-P and dorso-lateral axes using an electromagnet. After injection, the embryos are positioned above a permanent magnet and are let aged (2h to apply force during cellularization) in a wet chamber at 25°C.

### Calibration of the electro-magnet

To induce controlled forces we designed an electromagnet similarly to previous *in vitro* studies(Kollmannsberger and Fabry, 2007). A core of soft metal with a tip shape (Mumetal, Sekels Gmbh) is surrounded by 100 coils of copper cable alimented by a power supply in order to generate a solenoid. An additional radiator is placed in between the soft metal core and the cupper coils to evacuate the thermal dissipation. The electric current circulating within the cupper coils will generate a magnetic field that will be focused at the tip of the soft metal core. The magnetic force exerted on the paramagnetic micro-particle is directly proportional to the gradient of magnetic field(Kollmannsberger and Fabry, 2007). Therefore, the force would be the highest close to the tip and decay as a power law with the distance to the tip of the electro-magnet (Figure 1-figure supplement 1-B). In order to calibrate the magnetic force exerted on the micro-particle, we have established a calibration assay with micro-particle embedded in PDMS (Sigma Aldrich). Because its viscosity is well established, by measuring the velocity of the micro-particles within the magnetic field generated by the electro-magnet, one can determine precisely the force applied to the particle as a function of its distance to the tip for a specific current applied in the solenoid. For a current of 0.3A, the typical force-distance curve range from ~1nN at 60 μm from the tip to ~100 pN at 200 μm (Figure 1-figure supplement 1A-B). Due to geometrical constrains arising from the embryo shape, the typically used bead-magnet distances in our experiments were about 190μm (inset Figure 1-figure supplement 1B). This corresponds to forces of ~115pN (high force condition, for 0.3A) and ~50pN (low force condition, for 0.15A). Note that for currents above 0.5A the magnetic force saturates.

### Imaging and force application

Embryos were imaged at room temperature (22°C) using an Andor spinning disk confocal microscope. Z stacks of 4-5 sections spaced by 1 micron interval were collected every 5 seconds with 100X magnification.

The magnet was precisely positioned with a three-axis micromanipulator (Narishige UMM-3FC) mounted on the microscope stage. In the experiments, the electromagnet has been positioned approximately at 190 μm from the bead, and the force has been applied systematically for 65s. Data considered valid for analysis are cases when the bead is attached to the apical cell membrane. Occasionally, the bead detaches from the membrane, displaces in the basal direction or is attached to a wrong site in the cell for pulling experiment. In those cases, we did not include the data in the analysis.

### Analysis of the bead displacement

Analysis of the bead displacement was performed using Fiji. Tracking of the bead was performed on images acquired on the red channel where the auto-fluorescence of the particle is emitted. The following analysis step were implemented: (1) A median filter (radius 2) is applied on maximum intensity projections of each z stacks, (2) the projection is then manually thresholded, (3) MTrack2 plugin was used to automatically extract the x-y coordinate of the bead. In some occasions, samples showed drift. In order to detrend extracted bead displacements, we fit a linear function, *f* (*x*) = *mx* + *c*, to the bead displacement in the 100s preceding the force application. Assuming this drift pertained during the force application, we subtracted this linear trend from the bead displacement.

### Statistics

The table below reports a Welch test, checking the Null hypothesis that two sample have equal mean, without assuming they have the same standard deviation. All the conditions are compared with the respective WT at early and late cellularization for the data generated using the spring-dashpot model as reported in Figure2-figure supplement 1B.

|  | Condition | Stage | Delta | Parameter | cmax | cmin | p | p_values |
| --- | --- | --- | --- | --- | --- | --- | --- | --- |
| 0 | DMSO | Early Cell. | 43% | $K$ | 0.107667 | -0.061682 | - | 5,46E-01 |
| 1 | DMSO | Early Cell. | -12% | $\mu_1$ | 0.246043 | -0.611750 | - | 3,75E-01 |
| 2 | DMSO | Early Cell. | -36% | $1/\mu_2$ | 0.413671 | -0.505623 | - | 8,20E-01 |
| 3 | Colcemid | Early Cell. | 68% | $K$ | 0.071115 | 0.001740 | ** | 4,08E-02 |
| 4 | Colcemid | Early Cell. | 13% | $\mu_1$ | 0.564621 | -0.191551 | - | 3,16E-01 |
| 5 | Colcemid | Early Cell. | 134% | $1/\mu_2$ | 0.232951 | 0.101975 | ** | 3,37E-05 |
| 6 | RhoK | Early Cell. | 26% | $K$ | 0.038090 | -0.010224 | - | 2,43E-01 |
| 7 | RhoK | Early Cell. | 26% | $\mu_1$ | 0.734543 | 0.015826 | ** | 4,16E-02 |
| 8 | RhoK | Early Cell. | 6% | $1/\mu_2$ | 0.102569 | -0.086092 | - | 8,55E-01 |
| 9 | RhoK | Late Cell. | 14% | $K$ | 0.013819 | -0.007168 | - | 4,96E-01 |
| 10 | RhoK | Late Cell. | 13% | $\mu_1$ | 0.302732 | -0.139462 | - | 4,43E-01 |
| 11 | RhoK | Late Cell. | 0,09 | $1/\mu_2$ | 0.156488 | -0.258295 | - | 6,08E-01 |
| 12 | Low force | Early Cell. | 26% | $K$ | 0.062561 | -0.034499 | - | 5,21E-01 |
| 13 | Low force | Early Cell. | -4% | $\mu_1$ | 0.488387 | -0.621007 | - | 7,94E-01 |
| 14 | Low force | Early Cell. | 98% | $1/\mu_2$ | 0.380733 | -0.135324 | - | 2,93E-01 |
| 15 | Low force | Late Cell. | -26% | $K$ | 0.002841 | -0.014751 | - | 1,63E-01 |
| 16 | Low force | Late Cell. | -35% | $\mu_1$ | -0.017881 | -0.422326 | ** | 3,50E-02 |
| 17 | Low force | Late Cell. | -7% | $1/\mu_2$ | 0.404143 | -0.487061 | - | 8,33E-01 |

### Analysis of the experimental deformation field

Experimental images where aligned using the position of the bead at t=0s. Cell shapes were determined from the outlines of cell membranes at the onset and at the end of the period of force application, i.e. at t=0s and t=65s, respectively, using the software Packing Analyzer v.2.0(Aigouy et al., 2010). Cells where tracked using the same software. We determined the displacement vector, d(x), corresponding to the centroid movement between t=0s and t=65s of all cells within the field of view. Here, x=(x,y) is a vector denoting the position of the centroid of the cell at t=0s relative to the bead at t=0s. We considered the total set of obtained deformation vectors, {*d*(*x_k_*) *for k*=1, …, *M*}, as the readout of the experimental deformation field. Note that cells have a typical diameter of about ~5*μm*, which then defines the spatial resolution of the deformation field measurement.

### Vitelline envelope-apical cell surface distance estimation analysis

The estimation of the distance between the vitellin envelope and apical cell surface was performed using z-reslice along the AP axis (using FIJI) of z-stacks spaced by 0.5 μm of embryo expressing Resille-GFP injected with Dextran Texas-Red 70000MW(Molecular Probes) in the perivitelline space. We use the fluorescence intensity of Dextran Texas Red as a proxy for the position of the perivitelline space. With a homemade Matlab script, an average z-fluorescence profile is obtained by: 1) performing a sliding average of 5 pixels along the z-reslice and 2) by realigning in z each local average to the maximal dextran-RFP fluorescence intensity. A spline interpolation has been performed on the Resille-GFP intensity curves to determine the position of the maximum of intensity.

### Fly stocks

Stock used for live imaging: Resille-GFP (Morin et al., 2001) and Sqh^Ax3^, Sqh-GFP (Royou et al., 2004)

### Drug Microinjection

Developing embryos have been injected in early phase of cellularization using a Femtojet Injector (Eppendorf). Rho-K inhibitor (Y-27632 Sigma Aldrich) has been injected at 30mM; Colcemid (Santa Cruz Biotechnology) has been injected at 1mM and Dextran Texas-Red 70000MW(Molecular Probes) has been injected at concentrations between 7 and 25mg/ml. We estimate ~20 fold dilution in the embryo.

---

**Figure 1-figure supplement 1.**
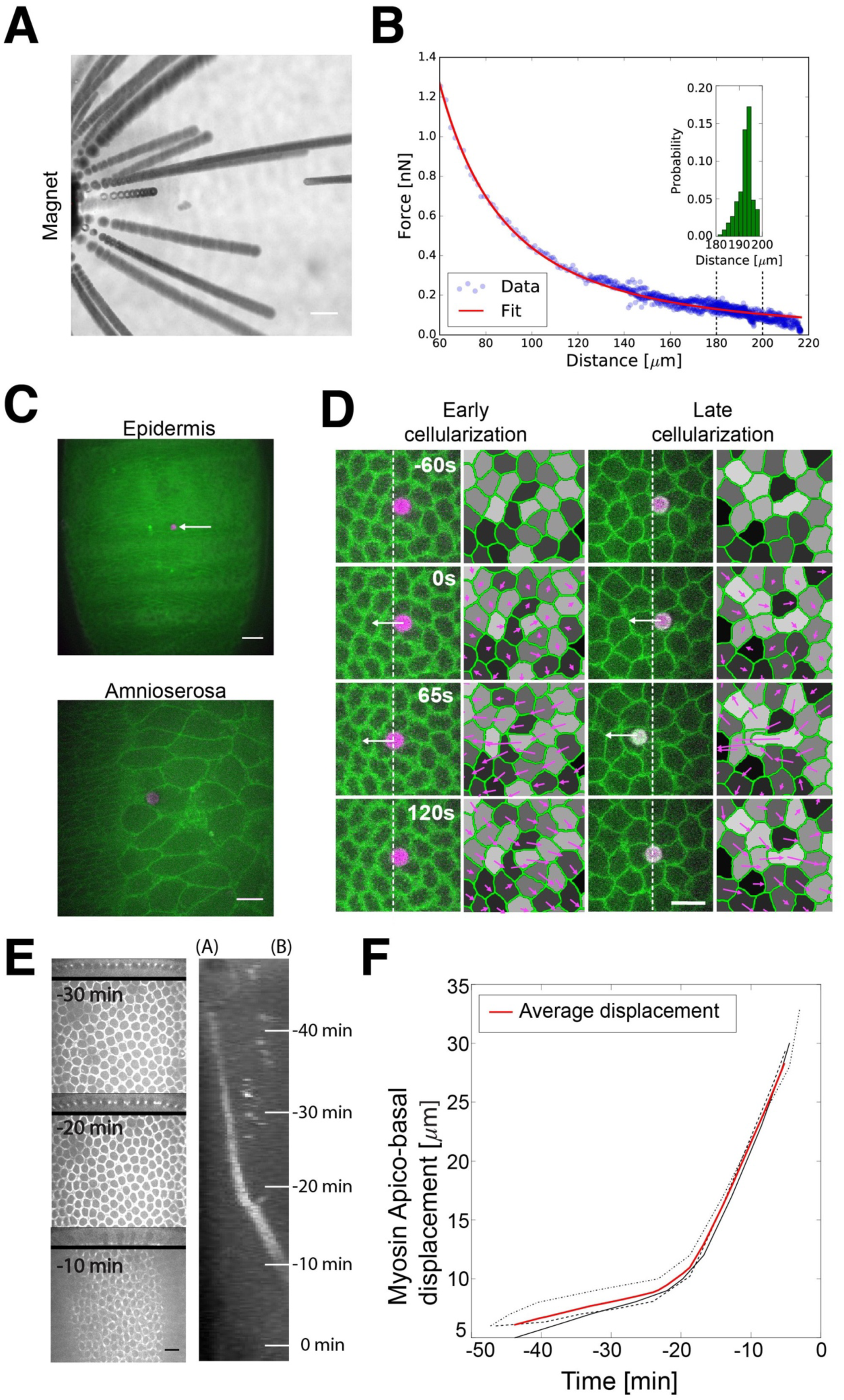
Force calibration, Bead positioning in tissues and cellularization process. (A) Time projection movie of 4.5 *μm* magnetic beads embedded in PDMS moving toward the magnet during application of the magnetic field generated with a current of 0.3A. The dark lines indicate bead trajectories. Scale bar is 20 *μm* (B) Example of force-distance calibration curves of 4.5 *μ*m beads (dots) and power law fit (in red). Inset shows the distribution probability of bead-magnet distances for the used force application experiments. (C) Snapshot of a single 4.5 *μm* bead (in magenta) embedded in the ventral epidermis (top, scale bar is 20 *μm*) and in the Amnioserosa (bottom, scale bar is 10 *μm*) at late stages of embryonic development. (D) Time lapse images showing the force application (high force condition) on a bead embedded into a single cell of a Resille-GFP embryo at early cellularization (on the left, >17 min before gastrulation) and late cellularization (on the right, <17 min before gastrulation) and the corresponding velocity fields. White arrows indicate when force is applied. White dashed lines indicate the left side of the bead at time -60s. Velocities are calculated between individual cell positions in the displayed time frames. Scale bar is 10 *μm* (E) Snapshot of Sqh-GFP embryo at -30, -20 and -10 min before the onset of gastrulation. Right: kymograph of myosin intensity along the z apico-basal direction of the tissue, integrated along the midline in the A-P axis, corresponding to the Sqh-GFP embryo shown on the left. A myosin intensity peak is displaced basally during the slow and fast phases of cellularization. Apical (A) is on the left and Basal (B) is on the right. The origin of time is set at the onset of gastrulation. Scale bar is 10 *μm* (F) Traces of the apico-basal displacement of the apicobasal intensity peak of myosin during cellularization in three different embryos (black lines), and their average (red line). The origin of time is set at the onset of gastrulation.

**Figure 2-figure supplement 1.**
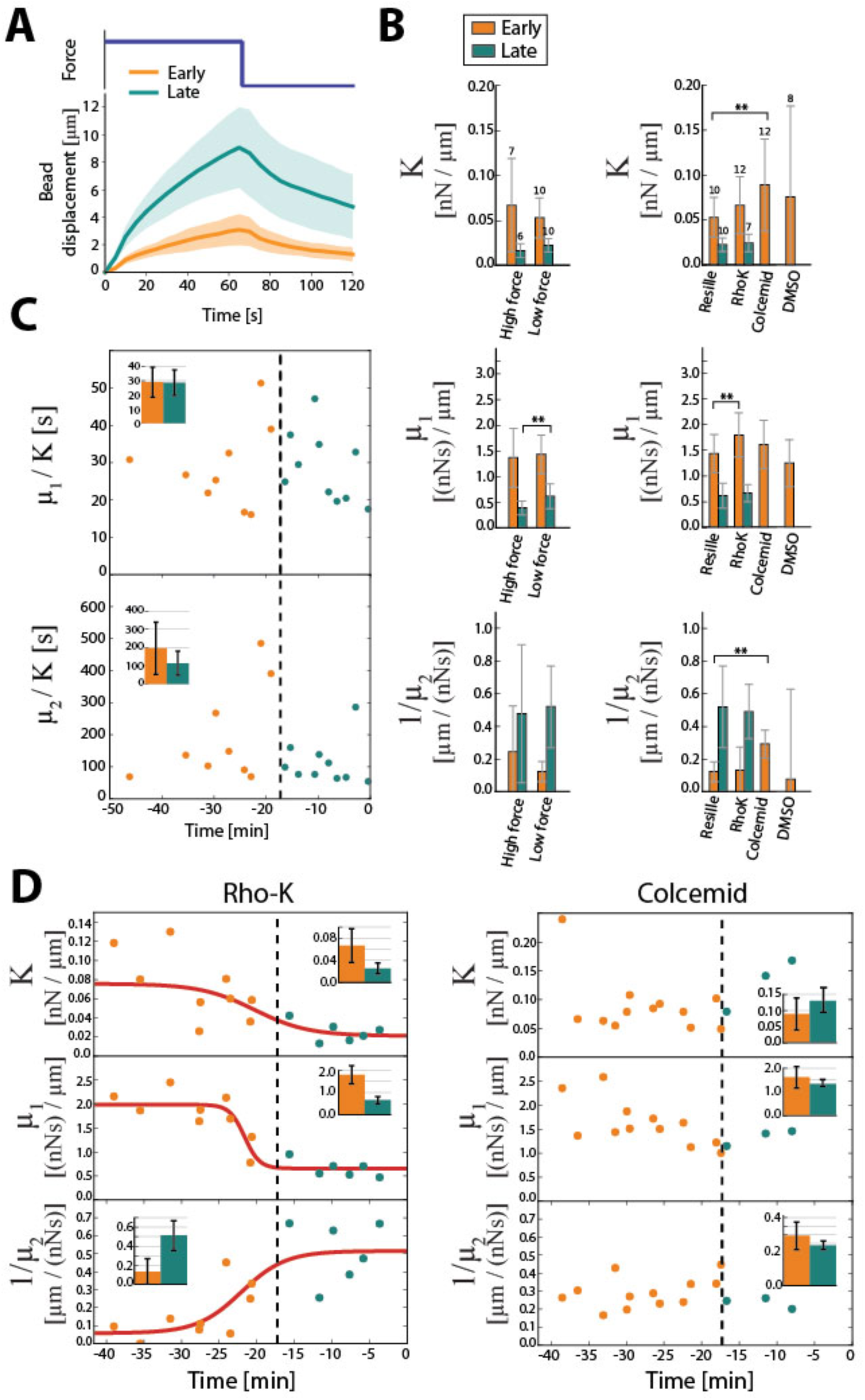
Analysis of bead displacements with a Maxwell Kelvin-Voigt model. (A) Average bead displacement for force steps of about 115 pN at early (n=10 applications) and late (n=10 applications) cellularization in Resille-GFP embryos. Shaded region represents standard deviations. (B) Estimation of stiffness *K*, viscosity coefficient *μ*_1_ and fluidity 1/*μ*_2_ using the Maxwell-Kelvin-Voigt spring-dashpot model. On the left, the parameters are estimated for WT embryo (Resille-GFP) at low force steps of ~50pN and high force steps of ~115pN. The estimated three parameters are approximately the same for 50pN and 115pN forces indicating that the tissue behaves linearly in this force regime. On the right, parameters are estimated for the same force amplitude (~115pN) for Resille-GFP expressing embryos and Resille-GFP embryos injected with Rho-K, Colcemid and DMSO. Numbers on the error bars are the number of force application. ** indicate p values smaller then 0.05 calculated using two sided t-test. Error bars represent standard deviations. (C) Viscoelastic timescales 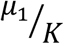 and 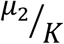 as a function of time. The origin of time is defined as the onset of gastrulation. The dashed lines indicate the separation between early and late cellularization. (D) Estimation of the three parameters stiffness *K*, viscosity coefficient *μ*_1_ and fluidity 1/*μ*_2_ over time for Resille-GFP embryos injected with Rho-K (left) inhibitor and Colcemid (right). For Rho-K injected embryos, we still observe a step-like behavior in the evolution of the three parameters. The red line is a sigmoidal fit recapitulating the time evolution of the parameters. For embryos treated with Colcemid there are no significant changes between early and late cellularization. In both cases the black dashed line represents the separation between early (in orange) and late (in blue) force applications. The insets show the mean value for each parameter for early and late cellularization. Error bars represent standard deviations.

**Figure 3-figure supplement 1.**
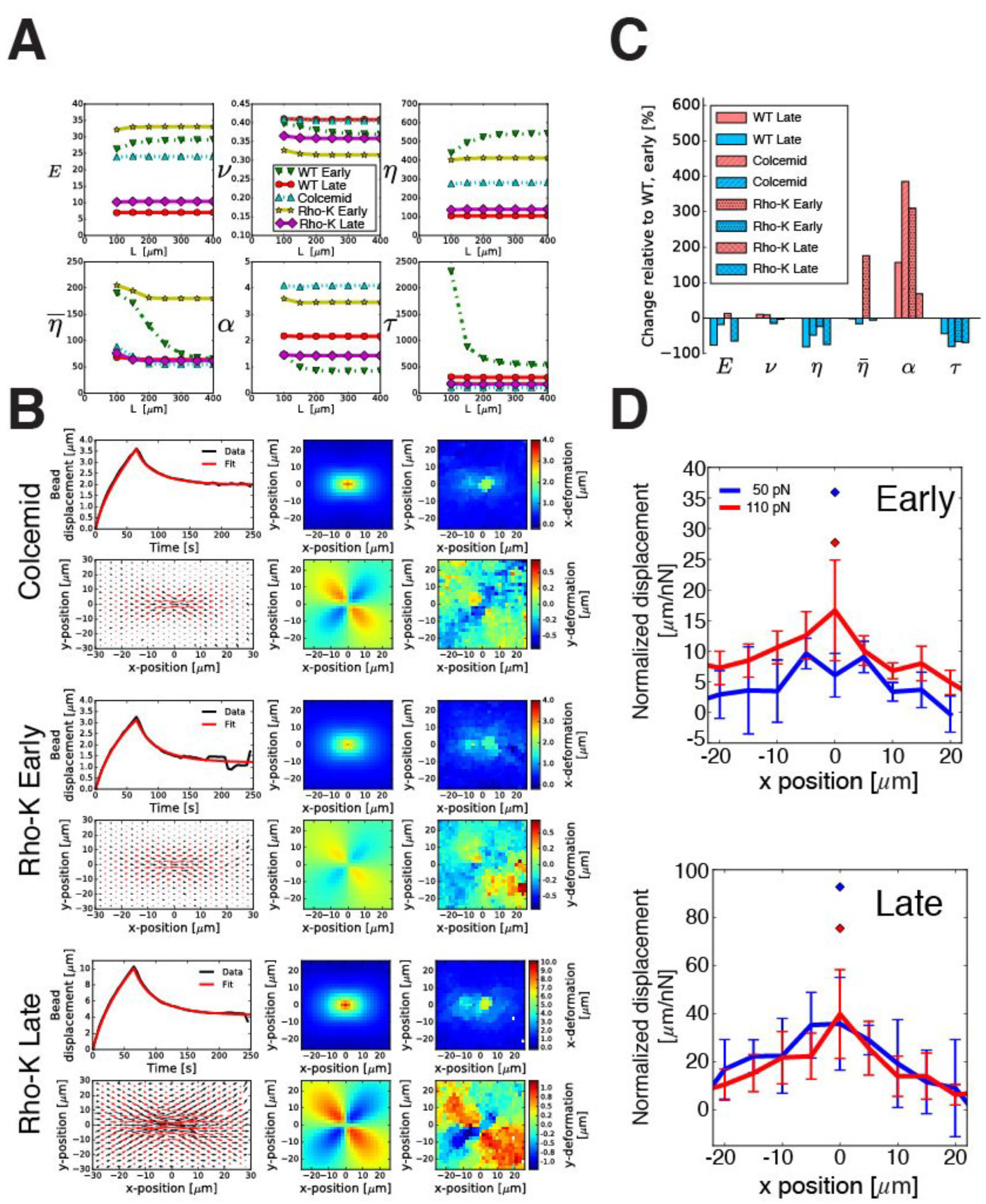
Analysis of bead displacement and deformation field with a 2D continuum mechanics model. (A) Mechanical parameters extracted from the fit of WT late cellularization bead displacement and deformation field as a function of the size L of the theoretical 2D viscoelastic tissue. Beyond 200μm, parameters are independent on the size of the 2D tissue. (B) Comparison of experimental and continuum description of average bead displacement in Colcemid (n=12), Rho-K Early (n=12) and Rho-K Late (n=7) and deformation field in Colcemid (n=8), Rho-K Early(n=8) and Rho-K Late (n=4). For all conditions, (top-left panel) average experimental and fitted bead displacement, (bottom-left panel) experimental (black arrows) and fitted deformation field (red arrows) and fitted and experimental x- and y-deformations (middle and right panels respectively) are shown. (C) Relative changes in the different mechanical parameters compared to WT early cellularization upon the different perturbations for early and late cellularization. (D) x-displacement of the tissues along the axis of force application, normalized by the pulling force magnitude taken at t=65s, for two force amplitudes 50 pN and 115 pN at early (Top) and late cellularization (Bottom). The Rhombuses indicates maximal bead displacements normalized by the force magnitude. Normalized displacements at 50pN and 115pN are similar, indicating a linear response of the tissue in this regime.

**Figure 3-figure supplement 2.**
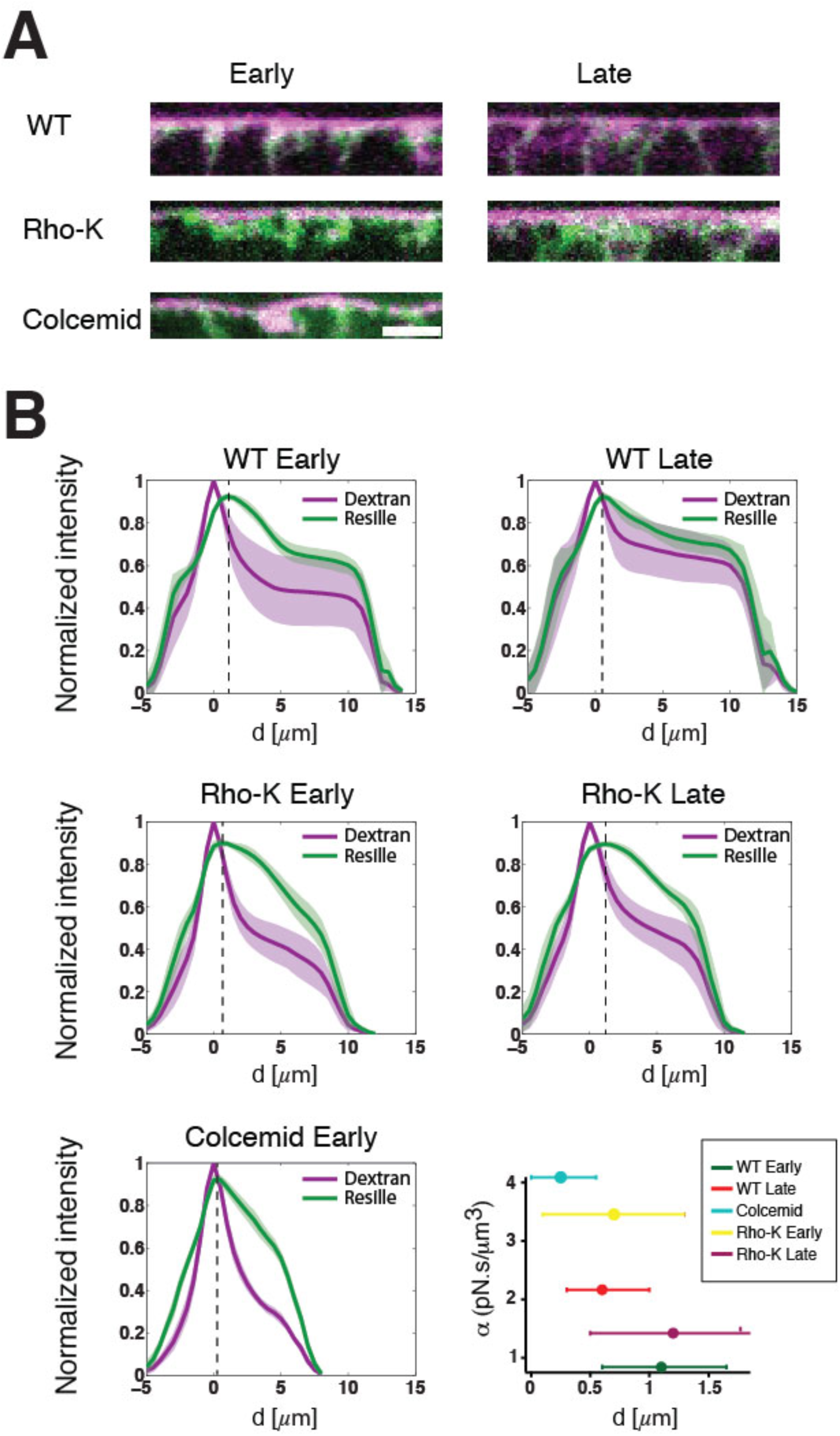
Estimation of perivitelline width for WT, Colcemid and Rho-K treated embryos. (A) Z-reslices along the A-P axis of the embryo showing dextran texas red (in magenta) injected in the perivitellin space and Resille GFP signal (in green) for WT embryos, Rho-K injected embryos at early and late cellularization and Colcemid injected embryos at early cellularization. Scale bar: 5μm (B) Average fluorescence intensity profiles of the dextran and membrane marker Resille-GFP labeling cellular membranes for WT and Rho-K treated embryos at early and late cellularization stages and in colcemid treated embryo at early stages (N_WTearly_=22 profiles on 3 embryos, N_WTlate_=21 profiles on 3 embryos, N_Rho-Kearly_=26 profiles on 3 embryos, N_Rho-Klate_=30 profiles on 3 embryos, N_Col_=10 profiles on 2 embryos). The dashed line defines the position of the maximum of the fluorescence peak for Resille (see methods). The bottom right graph shows the friction coefficient *α* (see fig 3-C) as a function of the dextran-Resille-GFP maximum peak intensity distance (as a proxy of perivitellin space width) for WT early and late and Rho-K inhibitor and Colcemid injected embryos. We can observe a linear relationship between friction coefficient and the inter-peak distance. The error bars represent the width of the Resille peak at 99% of the maximal height.

**Video 1 *Drosophila* larvae with a 4.5 *μ*m bead.**

Transmission movie of a drosophila larvae pre-injected at pre-blastoderm stage. The white arrow indicates the bead position.

**Video 2 Force application experiment at early cellularization.**

Time-lapse movie showing a force application experiment on a Resille-GFP embryo at early cellularization. The purple arrow indicates when a force of ~115pN is applied on the bead.

**Video 3 Force application experiment at late cellularization.**

Time-lapse movie showing a force application experiment on a Resille-GFP embryo at late cellularization. The purple arrow indicates when a force of ~115pN is applied to the bead.

**Video 4 Predicted tissue deformation upon force application at early cellularization.**

Kinetics of the deformation upon force application, resulting from the fit of pulling experiments at early cellularization with our 2D theoretical description. The left panel indicates the deformation field as a quiver plot, the middle panel the x-component and the right panel the y-component of the deformation field. The origin of time is the onset of the pulling force.

**Video 5 Predicted tissue deformation upon force application at late cellularization.**

Kinetics of the deformation upon force application, resulting from the fit of pulling experiments at late cellularization with our 2D theoretical description. The left panel indicates the deformation field as a quiver plot, the middle panel the x-component and the right panel the y-component of the deformation field. The origin of time is the onset of the pulling force.

**Video 6 Force application experiment at early cellularization in an embryo treated with Rho-K inhibitor.**

Time-lapse movie showing a force appllication experiment on a Resille-GFP embryo treated with Y27632 at early cellularization. The purple arrow indicates when a force of ~115pN is applied.

**Video 7 Force application experiment at late cellularization in an embryo treated with Rho-K inhibitor.**

Time-lapse movie showing a force application experiment on a Resille-GFP embryo treated with Y27632 at late cellularization. The purple arrow indicates when a force of ~115pN is applied.

**Video 8 Force application experiment at early cellularization in a Colcemid treated embryo.**

Time-lapse movie showing a force application experiment on a Resille-GFP embryo treated with Colcemid at early cellularization. The purple arrow indicates when a force of ~115pN is applied.

